# Endothelial tissue remodeling induced by intraluminal pressure enhances paracellular solute transport

**DOI:** 10.1101/2023.01.24.525266

**Authors:** Jean Cacheux, Aurélien Bancaud, Daniel Alcaide, Jun-ichi Suehiro, Yoshihiro Akimoto, Hiroyuki Sakurai, Yukiko T. Matsunaga

## Abstract

The endothelial layers of the microvasculature regulate the transport of solutes to the surrounding tissues. It remains unclear how this barrier function is affected by blood flow-induced intraluminal pressure. Using a 3D microvessel model, we compare the transport of macromolecules through endothelial tissues at mechanical rest or with intraluminal pressure, and correlate these data with electron microscopy of endothelial junctions. Upon application of an intraluminal pressure of 100 Pa, we demonstrate that the flow through the tissue increases by 135%. This increase is associated with a 25% expansion of microvessel diameter, which leads to tissue remodeling and thinning of the paracellular junctions. We recapitulate these data with the deformable monopore model, in which the increase in paracellular transport is explained by the augmentation of the diffusion rate across thinned junctions under mechanical stress. We therefore suggest that the deformation of microvasculatures contributes to regulate their barrier function.

---

Endothelial layers constitute physical barriers that act as gateways between the blood and the surrounding tissues^1,2^. They mediate exchanges of molecules, such as water, oxygen or proteins, and hence contribute to tissue homeostasis. Dysregulation of the barrier function contributes to the pathophysiology of many diseases, including cancer and infections. Endothelial tissues also constitute physical barriers that have to be crossed for efficient delivery of therapeutic agents^3^. The passage of solutes across these barriers occurs via two non-mutually exclusive mechanisms of transcellular and paracellular transport^4^. The transcellular route crosses apical and basal cell membranes and is mostly mediated by transcytosis, an energy-dependent trafficking of vesicles across the endothelium^5^, or the dynamic formation of patent openings^6^. In contrast, the paracellular pathway, which passes through intercellular spaces between contacting cells^7^, is considered to be passive. It is maintained by the interendothelial junctions, which are mediated by tight junction proteins and adherens junction proteins^8,9^, and/or by rearrangement of their architecture. While the relative strength and size selectivity of paracellular *vs*. transcellular pathways is tissue-specific and remains the subject of intense research *in vivo*^10–12^, it is considered that, except for cerebral capillaries in which tight junctions are most abundant^13^, small hydrophilic molecules transit through the paracellular route in other microvasculatures^14–16^.

In 2D cell culture systems, the paracellular transport pathway is most frequently characterized by trans-endothelial electrical resistance (TEER)^17–19^. This technique, which consists in recording the ionic current across a cell tissue upon application of an electric field, provides a readout that correlates with the structural integrity of endothelial barriers. Paracellular transport can also be assayed by the leakage or macromolecular assay (MA), which consists in tracking the temporal^20–23^ or spatial^24,25^ redistribution of fluorescently-labeled hydrophilic dextran through the tissue^26^. The readout of the MA is most frequently the diffusive permeability, which can be used to model the intercellular spaces as pores of characteristic diameter ∼5 nm for mesenteric capillaries^7^. This estimate is however obtained in the absence of intraluminal pressure, and the consequences of this constitutive cue of the vascular system on transport across the endothelium remain unclear^27^. The pressure gradient across the barrier is indeed expected to create a flow in the intercellular gaps, and in turn to enhance the apical to basal flux. However, whether diffusion or convection is the dominant transport mechanism in the paracellular pathway is debated in the literature^28^.

In this study, we set up a tissue-engineering approach to compare the strength of paracellular transport in endothelial tissues with or without intraluminal pressure. Using the transport at mechanical rest as a reference, we prove that intraluminal pressure enhances paracellular flux by 135%, strains the tissue by 25%, induces reorganization of the cytoskeleton associated actin stress fiber formation, and remodels the tissue though the formation of clusters of endothelial cells. Electron microscopic observation of intercellular junctions shows that the thickness of the paracellular junctions decreases by ∼ 50% in the stretched regions of the tissue. We show that the conventional monopore model^16^ of paracellular transport, in which the deformation of the tissue induced by intraluminal pressure is ignored, fails to account for this data. We thus propose the deformable monopore model (DMM) to integrate the consequences of the morphological change of paracellular junctions under pressure on barrier function properties.

## Experiments and models

### Principle of the static and pressure assays

We fabricated microvessels (MV) in collagen gels of 200 μm in diameter and 6 mm in length (Fig. 1A-B and Methods) following the protocols described in ref. ^29,30^. After two days of maturation, we placed a 3D printed device on top of the MV chip, which enabled us to control and maintain the level of liquid stably over time (Fig. 1C), and in turn to set the conditions of transport across the endothelial barrier. Specifically, the level of liquid over the two inlets of the MV and the collagen matrix was even in the static assay (left panel in Fig. 1D-E). We thus insured a constant pressure on the basal and apical sides of the tissue, and the passage of fluorescent macromolecules across the tissue was only forced by diffusion. In the pressure assay (right panel in Fig. 1D-E), we did not fill the central reservoir on top of the collagen gel with liquid in order to apply an intraluminal pressure of 100 Pa (1 cm of fluid height difference). The fluid height difference then remained constant in the time course of pressure assay experiments because the permeation flow through the MV barrier was too low to change the volume in the inlet reservoirs.

**Figure 1:**
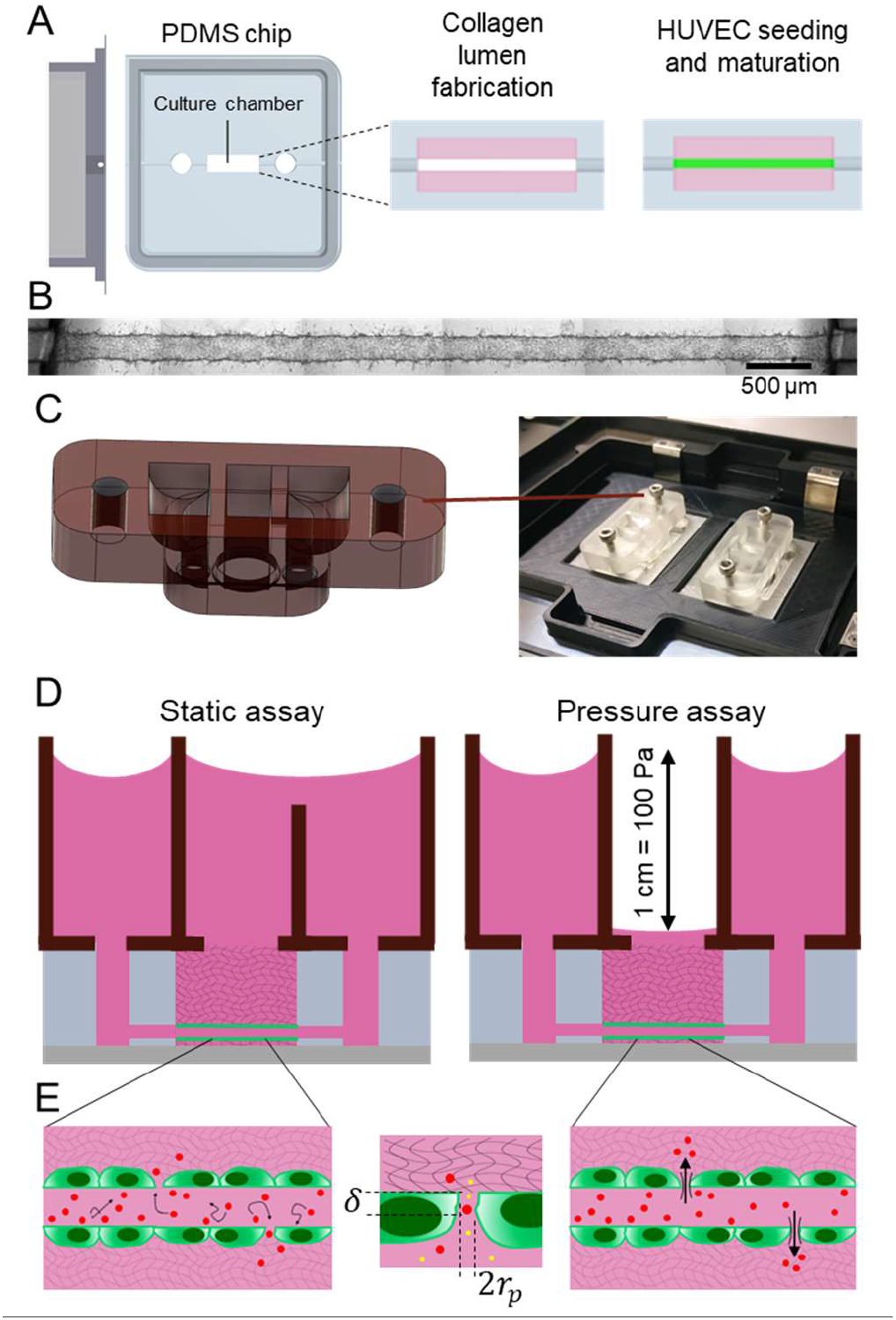
MV fabrication and characterization by the static and pressure assays. **(A)** Representation of the microvessel chip and the consecutive fabrication steps starting from Human Vascular Endothelial Cells (HUVEC). The chip is represented in gray, the collagen gel in pink, and the endothelial tissue in green. **(B)** Microscopic view of a MV after two days of culture. The scale bar corresponds to 500 µm. **(C)** Layout of the 3D printed device for the static assay, and photograph of the device in operation on an inverted microscope. **(D)** Schematic representation of the devices to run the static and pressure assays. Intraluminal pressure is produced by 1 cm of hydrostatic pressure. **(E)** The left and right panels represent the transport of tracers through the barrier using diffusion or pressure as actuation scheme, respectively. The central panel shows the monopore model, as defined by the pore radius r_p_, density n, and thickness δ. We investigate the passage of two macromolecules with different sizes across paracellular junctions, as shown with red and yellow circles.

### Models of the static and pressure assays

The static and pressure assays started by the injection of two dextran dyes in the lumen at *t* = 0. Their spatial redistribution was then monitored by confocal microscopy in the equatorial plane of the MV. Assuming an axisymmetric geometry, the dynamics of the concentration *C*(*r, t*) of macromolecules in the collagen matrix with *r* the distance to the center of the lumen and *t* the time is governed by Fick’s second law of diffusion with convection:

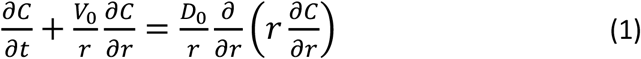

with *D*_0_ the tracer diffusion coefficient in collagen gels and *V*_0_ the permeation flow velocity across the barrier (*V*_0_ is null in the diffusion assay). The endothelial barrier sets the flux of macromolecules per unit of tube length that leaks out from the lumen to the collagen. In the static assay, this flux *J_D_*(*t*) is determined by the difference between the apical and basal concentrations *C*_*in*_ and *C*_*out*_, respectively, multiplied by the diffusive permeability *ℒ_D_*^16,31^:

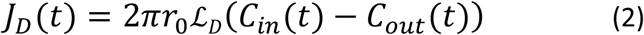

with *r*_0_ the radius of the tube.

Convective and diffusive solute transport contribute to the flux *J_p_*(*t*) in the pressure assay. It can be expressed with the Patlak equation^7^:

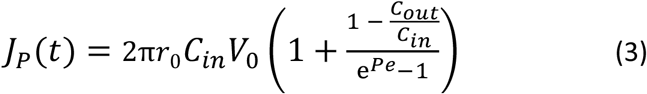

with *Pe* = *V*_0_/*ℒ_D_* the Peclet number that represents the ratio of the convection to diffusion rate in the intercellular gaps. On the vanishing *Pe* limit, Eq. (3) is the same as Eq. (2). Note that Eq. (3) is readily obtained by computing the flux through a single pore^32^, and summing the total flux over all the pores traversing the tissue. In the following, we describe and check the consistency of two methods to infer *ℒ_D_* and *V*_0_/*ℒ_D_*.

### Extraction of ℒ_D_ from the static assay

The flux *J_D_*(*t*) across the barrier in the static assay is equal to the temporal variation of the number of macromolecules *N*_*out*_(*t*) in the collagen gel. This number *N*_*out*_(*t*) can readily be extracted from confocal images by computing the integral of the concentration profile 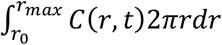 between the MV basal layer to the limit of the field of observation *r*_max_ (i.e., along the yellow dashed line in Fig. 2A). Hence, the diffusive permeability can be expressed from Eq. (2):

**Figure 2:**
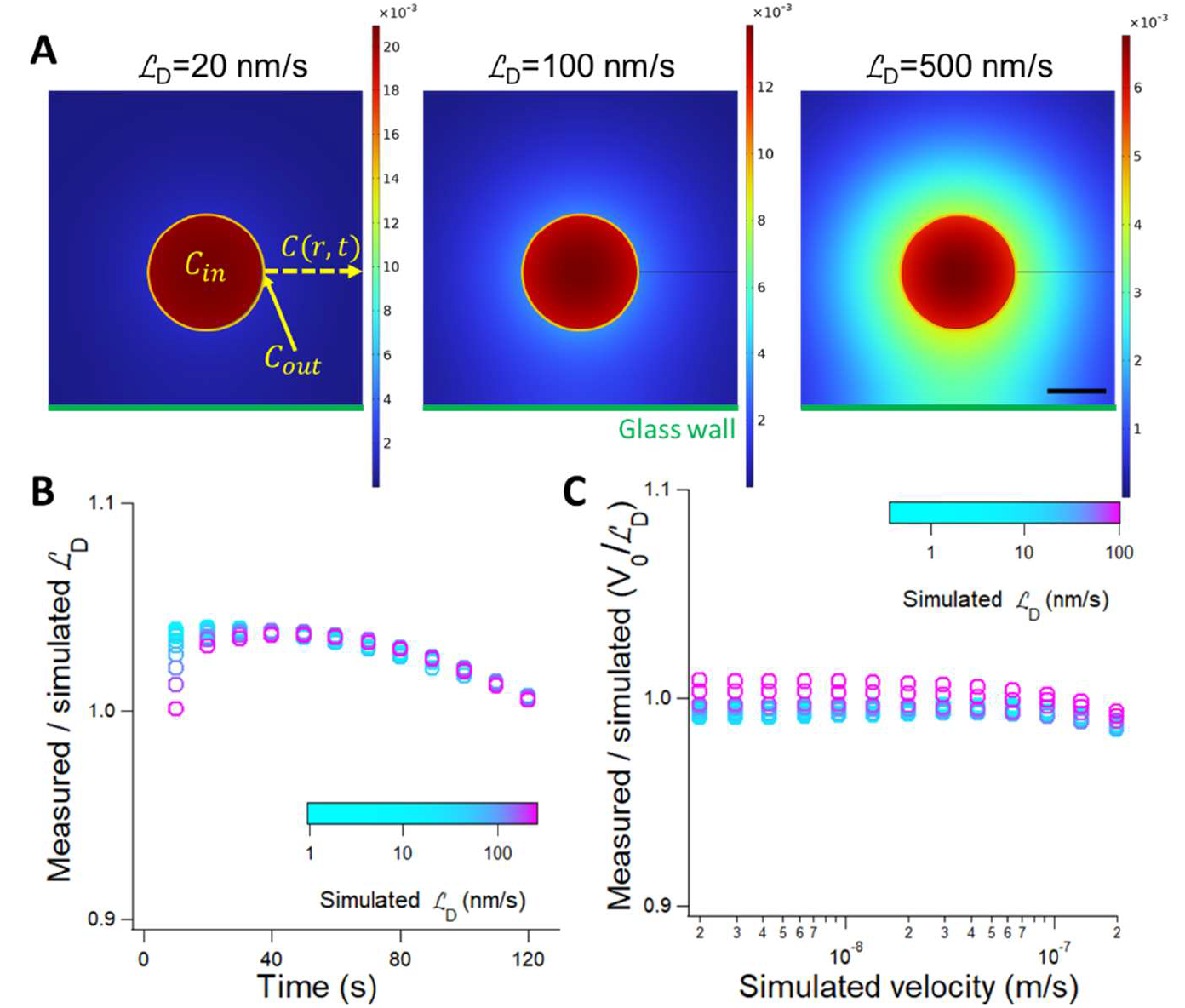
Validation of the static and pressure assays by finite element modeling. **(A)** The snapshots represent the cross-section of the MV with the lumen shown in red, the cell layer in orange, and the collagen gel in blue. In the static assay, the tissue is modelled by its diffusive permeability ℒ_D_, as indicated in the legend, and the simulations represent the concentration fields after a time lag of 200 s (the three heat maps are in units of mol/m^3^). We extract the basal and apical concentrations C_out_ and C_in_, respectively, and the concentration profile C(r, t) along the dashed yellow arrow. The scale bar corresponds to 100 µm. **(B)** The plot reports the ratio of the measured to the simulated ℒ_D_, as indicated in the color scale, as a function of time. **(C)** The plot presents the ratio of the measured V_0_/ℒ_D_ divided by this input parameter of the simulations as a function of the permeation velocity across the tissue V_0_. Data points are color-coded according to the value of the diffusive permeability, as indicated in the color scale.

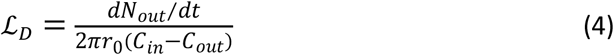

We validated the accuracy of Eq. (4) by running finite element simulations (see Methods) in order to take the geometrical settings of the experiments into account. We extracted *C*_*in*_, *C*_*out*_, and *dN*_*out*_/*dt* from the simulations (Fig. 2A), determined *ℒ_D_* using Eq. (4) and compared this parameter to the input of the simulations (see legend of Fig. 2B). We concluded that this method was accurate because the difference between the measured and the simulated *ℒ_D_* was less than 3% during the first 200 s of the simulation (Fig. 2B). Notably, the smooth decrease of *ℒ_D_* over time is readily explained by the escape of tracers from the field of observation of the microscope, which biases the determination of the flux *J_D_*(*t*) due to the error on the integral of *C*(*r, t*).

### Extraction of V_0_/ℒ_D_ from the pressure assay

For the analysis of the pressure assay, one should compare the rate of convection and diffusion in the collagen gel, which is gauged by the ratio *V*_0_*r*_0_/*D*_0_. Taking typical values of *r*_0_, *V*_0_, and *D*_0_ of 100 µm, 0.1 µm/s, and 250 µm^2^/s (see below), the latter ratio is ∼0.05. Hence, the convection rate is negligible compared to the diffusion rate. It implies that the equation governing fluorescence spatial redistribution in the pressure assay is the same as in the static assay, but the flux of molecules crossing the barrier *J_p_*(*t*) is different. By computing the ratio *J_p_*(*t*)/*J_D_*(*t*), we obtain an expression of *V*_0_/*ℒ_D_*:

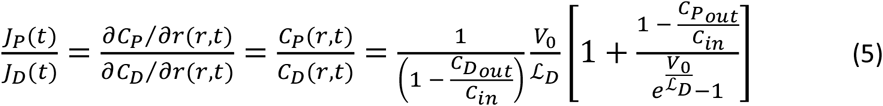

with 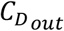 and 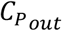 the basal concentrations in the static and pressure assays, respectively. Consequently, *V*_0_/*ℒ_D_* can be inferred from the numerical inversion of Eq. (5) knowing the concentration profile measured with the pressure and static assay.

We validated this method using finite element simulations in which the permeation velocity *V*_0_ and the diffusive permeability *ℒ_D_* varied in the range of 1 to 200 nm/s. For each simulation, we extracted the concentration profile for the pressure assay after 30 s and normalized it with the output of simulations of the static assay in order to measure *V*_0_/*ℒ_D_* using Eq. (5). We computed the ratio of the resulting value of *V*_0_/*ℒ_D_* divided by the input of the simulations, and plotted this parameter in Fig. 2C. This graph showed that the measured value of *V*_0_/*ℒ_D_* was consistent with that of the simulation, and the error was lower than 5% for a broad range of experimental conditions.

### Predictions of the monopore model of paracellular transport for ℒ_D_ and V_0_

Our goal is to relate the diffusive permeability and permeation velocity to the structure of paracellular pores. The pore model^16^ is defined by the pore radius *r_p_*, the density of pores per unit of surface *n*, and the barrier thickness *δ* (see schematics in the middle panel of Fig. 1E). The tissue porosity *φ* is defined by the area of void space over total surface:

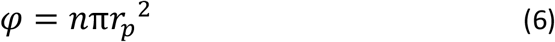

According to Fick’s law, *ℒ_D_* is dictated by the gradient of dextran concentration from the basal to apical side of the tissue. Assuming that this concentration gradient is constant across the pore (a common approximation in separation science^33^), we deduce a relationship between *ℒ_D_* and the geometry of the pores

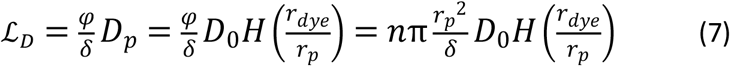

with *D_p_* the diffusion coefficient in the pore. The diffusion coefficient *D_p_* is dependent on the size of tracers, as described by the hindrance factor *H*(*r_dye_* /*r_p_*)^32^.

The ease of fluid flow through a barrier is described by the permeability of the endothelial tissue κ, which relates the permeation flow velocity to the pressure gradient across the barrier thickness. The permeability of an array of parallel pores is^34^

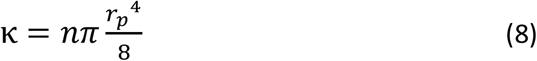

Combining Eq. (6) and (8), we obtain a well-known relationship of the form κ = *φr*_*p*_^2^/8 for porous media^35^. Provided that the cell layer is much more impermeable than the surrounding collagen scaffold (as validated by simulations, not shown), we posit that *V*_0_ is proportional to κ, and derive a relationship between *V*_0_ and *ℒ*_*D*_ using Eq. (7):

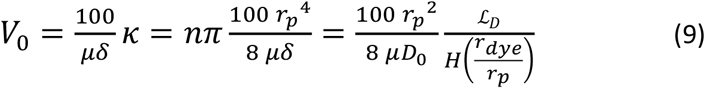

According to Eq. (9), the ratio *V*_0_/*ℒ_D_* is a function the pore radius independently of the density of pores and the barrier thickness.

### Deformable monopore model

Intraluminal pressure strains the tissue and remodels the structure of MV (see more below). This reorganization is associated to a reduction of the barrier thickness, and to an onset of the diffusive permeability, that we denote 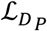. Integrating this term in Eq. (3), we reformulate Eq. (5) with the deformable monopore model (DMM) to include the consequences of the deformation of the MV induced by intraluminal pressure:

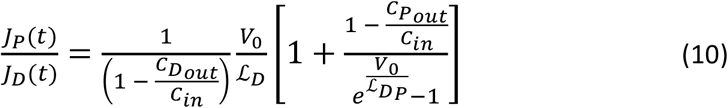

## Results and discussion

### The static assay predicts a size of paracellular pores of 24 nm

We first proceeded with the diffusion assay to characterize the barrier properties of MVs, because its operation requires the same conditions as those of the maturation of the tissue (*i*.*e*., without intraluminal pressure). We used nine MVs and ten fixed MVs, which were characterized by an average radius *r*_0_ of 102 +/- 2 µm (see Supplementary Fig. S1 for edge detection strategy) that was comparable to the radius of the lumen before cell seeding of 100 µm. The static assay was operated with two dextran probes of 4 and 70 kDa simultaneously injected in the lumen of MVs. The spreading of these macromolecules was then monitored by confocal microscopy for ∼200 s (Fig. 3A and Methods). Following the method based on Eq. (4), we measured the diffusive permeability by extracting the intraluminal and basal concentrations *C*_*in*_ and *C*_*out*_ (green and orange rectangles in Fig. 3A, respectively), and the number of tracers *N*_*out*_ in the collagen matrix by spatial integration of the intensity profile in the red dashed area in Fig. 3A. In Fig. 3B, we report the variation of *C*_*in*_ and *C*_*out*_ over time, showing opposite trends associated to a decrease and an increase, respectively (green and orange datasets), because the dextran molecules escape from the lumen by diffusion. The steepness of the concentration gradient across the barrier thus decreases, and the cross-barrier flux *J_D_*(*t*) is expected to decrease (see Eq. (2)). This trend is readily confirmed by the softening of the slope of *N*_*out*_ over time (red dashed curve in Fig. 3B). We use these different readouts to extract the diffusive permeability for the 4 and 70 kDa dextran (Fig. 3C), which appeared to be roughly constant over time. Note that the diffusive permeability was estimated on both sides of the MV at each time step of the time series. We computed the temporal average of the diffusive permeability of 91.3 +/- 8.5 and 10.9 +/- 2.2 nm/s for the 4 and 70 kDa dextran, respectively (purple and orange datasets in Fig. 3C).

**Figure 3:**
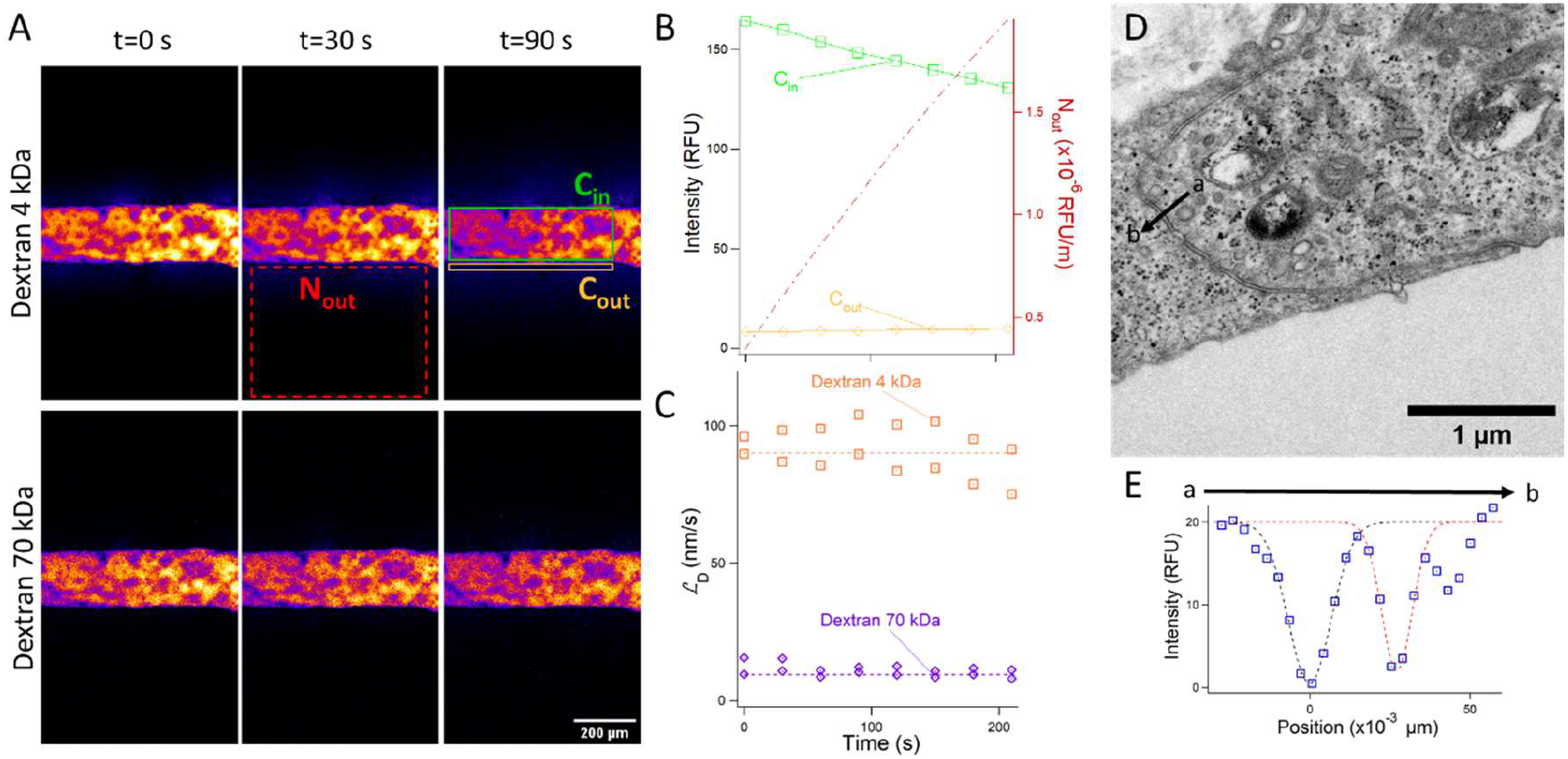
MV characterized by the static assay. **(A)** The two confocal time series represent fluorescence spatial redistribution for the 4 and 70 kDa dextran. The number of molecules N_out_ that crossed the barrier is measured at each time step in the collagen matrix (red rectangle). The intraluminal and basal concentrations C_in_ and C_out_ are inferred from the maximum and mean intensity in the green and yellow rectangles, respectively. **(B)** The graph shows the temporal variation of C_in_ and C_out_ on the left axis for the 4 kDa dextran as well as and N_out_(t) on the right axis with the color code of panel A. **(C)** The plot presents the diffusive permeability for the 4 and 70 kDa dextran as a function of time. At each time point, the diffusive permeability is measured on both sides of the MV. Dashed lines correspond to temporal averages. **(D)** Transmission electron micrograph of one endothelial cell junction. **(E)** The plot shows the spatial variation of the signal along the black arrow in (D). Dashed lines are Gaussian fits to measure the size of the paracellular gap.

This operation was repeated on the nine MV samples to estimate the average diffusive permeability *ℒ_D_* of 87 +/- 13 nm/s for the 4 kDa dextran, which was comparable to the estimate in *in vitro* systems^36^, but was slightly higher than the value reported for skeletal muscle microvasculatures of 10 nm/s^37^. *ℒ_D_* was 14 +/- 3 nm/s for the 70 kDa dextran (*i*.*e*., 6.2 times lower than for the 4 kDa dextran). We performed the same characterization with the ten fixed samples, and obtained very comparable values for the diffusive permeability of 107 +/- 20 nm/s and 14 +/- 3 nm/s for the 4 kDa and 70 kDa dextran, respectively, suggesting that the structural integrity of the tissue was marginally altered by the fixation process.

The variation of the diffusive permeability of the 4 *vs*. 70 kDa dextran is 6.2-fold, whereas the change in diffusion coefficient between these macromolecules is 3.9-fold (see characterization in Supplementary Fig. S2). This difference is explained by the confinement of the tracers in the intercellular gaps (see the scheme with red and yellow probes in Fig. 1D). Confinement is indeed known to slow down the diffusion of probes across pores (see Eq. (7)), and this consequence increases as the size of the tracer *r_dye_* becomes comparable to that of the pore *r_p_* ^32^. The variation of the diffusive permeability enables us to fit the level of confinement *r_dye_* /*r_p_*, and in turn to infer the size of paracellular pores. This approach can be performed on the nine MVs independently in order to obtain an average pore size *r_p_* of 24 +/- 3 nm (note that the fitted pore size is the same for fixed MVs). The analysis is strongly supported by transmission electron microscopic observation of MV thin sections (Fig. 3D and Methods), which show a pore radius of 13 +/- 2 nm based on an average over 4 samples (Fig. 3E). Consequently, the structure of paracellular gaps inferred from functional diffusion-based measurements are corroborated by electron microscopic inspection, implying that transport data provide a direct readout of the integrity of endothelial tissues.

### The pressure assay predicts a size of paracellular pores of 179 nm

Whether the structure of paracellular pores is altered by intraluminal pressure remains unclear. Hence, we performed the pressure assay on the same MV samples in order to obtain a quantitative comparison of the paracellular flux with or without stress on the tissue. We set the intraluminal pressure to 100 Pa in order to apply a mechanical stress comparable to the pulsatile pressure in human capillaries of ∼500 Pa^38^, and injected the 4 kDa dextran in the lumen (Fig. 4A). We extracted the concentration profile after a time lag of 30 s (see model section and red dataset in Fig. 4B). This time interval was selected to obtain a sharp concentration profile easily comparable to the outcome of the static assay (black dataset in Fig. 4B). The greater amplitude of the concentration profile in the pressure assay indicated that intraluminal pressure increased the transport rate across the barrier. Upon normalization of the concentration profile, the responses of the pressure and static assays were superposed (pink dashed lines *vs*. black datasets in Fig. 4B). This response directly showed that fluorescence redistribution in collagen gels was dominated by diffusion even with intraluminal pressure, as explained in the model section. This statement was further borne out upon characterizing one leaky barrier, which was obtained by loading a low number of cells in the duct, because the concentration profile in the pressure assay did not match that of the static assay (Supplementary Fig. S3). The ratio of the cross-barrier flux in the pressure *vs*. static assay *J_p_*/*J_D_* was then averaged over the nine samples, showing an onset of 135 +/- 43% (red dataset in Fig. 4C). In the ten fixed samples, this ratio was comparatively lower and equal to 67 +/- 37% (blue dataset in Fig. 4C).

**Figure 4:**
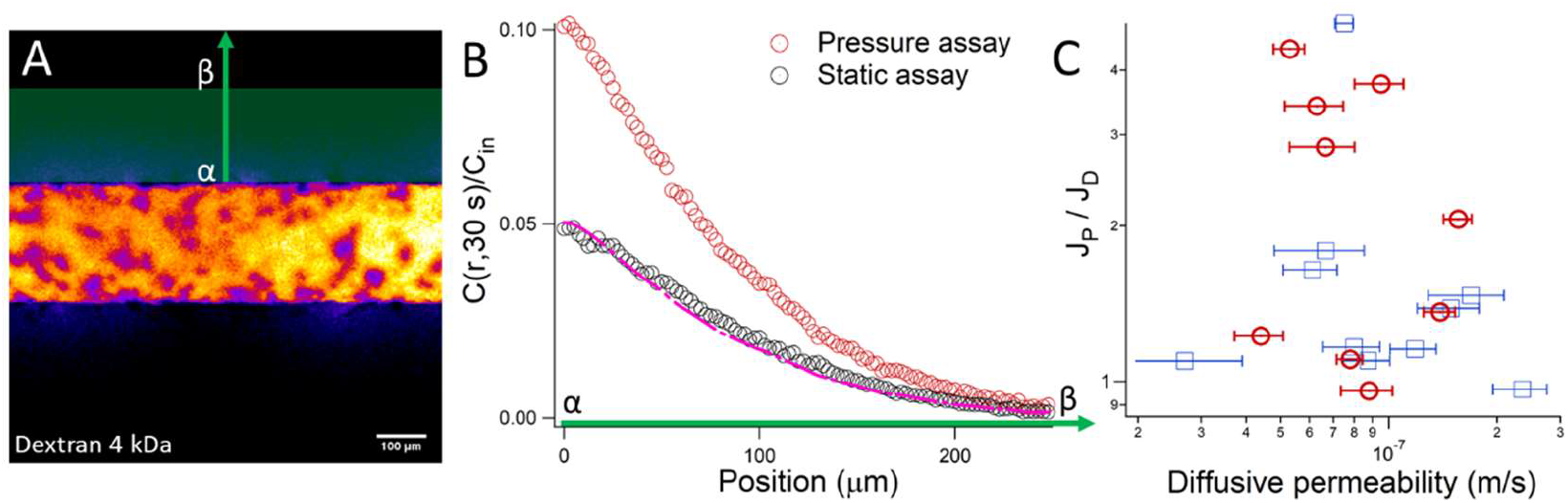
MV characterized by the pressure assay. **(A)** Fluorescence micrograph of one MV injected with the 4 kDa dextran after 30 s of acquisition. **(B)** The red and black datasets represent the concentration profile as obtained from the pressure and static assay, respectively. The profiles are recorded along the green axis in (A) between the marks α and β. The pink dashed line is response of the pressure assay after normalization of its maximum to that of the static assay. **(C)** Ratio of the flux across the MV barrier in the pressure to static assay as a function of the diffusive permeability ℒ_D_ for live and fixed samples (red and blue datasets, respectively).

According to Eq. (5), the ratio *J_p_*/*J_D_* enables us to determine the ratio *V*_0_/*ℒ_D_* of the permeation flow velocity to the diffusive permeability, which was equal to 1.91 +/- 0.54. The fact that this factor is greater than 1 indicates that the application of intraluminal pressure creates a flow in the paracellular pathway that conveys fluorescent macromolecules more rapidly than the diffusion process alone. Further, we use the predictions of the pore model (Eq. (9)) to determine the paracellular pore size from *V*_0_/*ℒ_D_* (note that the 4 kDa dextran is much smaller than the pore size, implying that the hindrance factor is equal to 1). We deduce that the average paracellular pore size *r_p_* of 179 nm is 7.5-fold wider than the dimension inferred from the static assay of 24 nm. The integrity of MVs hence appears to be strongly altered by the application of intraluminal pressure, prompting us to investigate their structure by transmission electron microscopy.

### Structural analysis of MV in the pressure assay

Before the fixation process, we first noted that the radius of MVs increased by 25 +/- 1% from 102 +/- 2 to 128 +/- 2 µm and was stable over time (not shown). This level of deformation has been shown to activate endothelial remodeling^39,40^, leading us to inspect the structure of the cytoskeleton, the morphology of the tissue and the geometry of intercellular gaps. MVs were fixed immediately after the pressure assay, and we started the analysis at the largest scale using optical microscopy and labeling thin sections of the tissue with toluidine blue^41^ (Fig. 5A-B). The samples of the static assay showed an even distribution of elongated cells with flat nuclei over the contour of the vessel, as expected for endothelial tissues (arrow in Fig. 5A). The application of intraluminal pressure induced a drastic morphological change associated to the formation of clusters of cells (arrowheads in Fig. 5B). These clusters represented a fraction of the contour of the tissue of 35.4% that was 4.4 times larger than in the static assay of 8.1%. Focusing on the cellular level, we then performed immunostaining of the actin cytoskeleton, which showed predominant cortical actin rings patterns at rest whereas transcellular actin stress fibers were frequent with intraluminal pressure (Fig. 5C-D), in agreement with experiments of cyclic stretch on 2D endothelial tissues^40^. Given that actin mediates the movement of nuclei within the cell^42^, we suggest that the formation of stress fibers, which is associated to tensile forces^43^, accounts for the clustering of cell nuclei. Despite the remodeling of the tissue, the patterns of vascular endothelial cadherin (VE-Cad), which has a key role in the maintenance of vascular integrity^2^, were not altered by the mechanical stimulation (Fig. 5C-D). Notably, this observation was not comparable to the readout of cyclic stretch experiments, which showed the partial disassembly of adherens junctions associated to barrier function weakening^39^.

**Figure 5:**
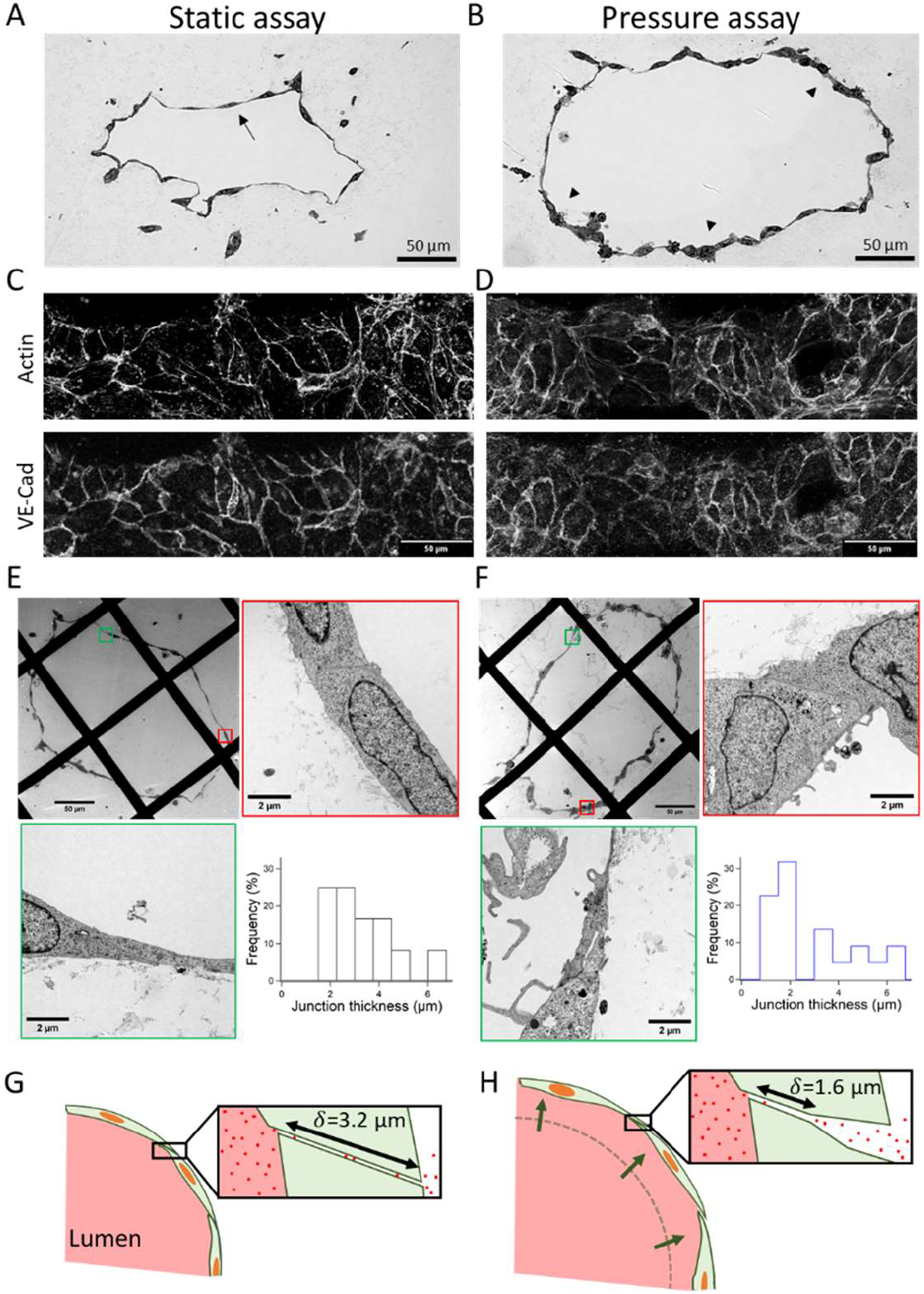
Structure of MVs in the static and pressure assay. The results of the static assay are shown in the panels **(A, C, E)**, and those of the pressure assay in **(B, D, F). (A-B)** Optical micrographs of MV sections stained with toluidine blue. The arrow in (A) shows the lining of endothelial cells, and the arrowheads in (B) the clusters of endothelial cells. **(C-D)** Maximum intensity projection of confocal micrographs obtained by staining MV with phalloidin for the detection of fibril actin and antibody for VE-Cad. The confocal stacks are recorded at the bottom of the MV. **(E-F)** Transmission electron micrographs of MVs at different levels of magnification. The red and green outlines correspond to the zooms of the squares in the low magnification image. The histograms show the distribution of paracellular junction thickness with 12 counts in (E) and 22 in (F). **(G-H)** Representation of the deformable monopore model to account for the data from the static and pressure assays.

We finally aimed at clarifying whether the structure of the paracellular junctions was altered by the mechanical stress using high magnification transverse electron microscopy. At mechanical rest, we noted two different patterns of junctions (Fig. 5E). In contacting cells with close nuclei, the paracellular cleft across the endothelium was mainly aligned along the apical-to-basal axis, and the thickness of the junction was commensurate with that of the nuclei on the order of 3 to 5 µm (micrograph with the red contour in Fig. 5E). In the thin regions of the tissue, most junctions were tilted with respected to the basal to apical direction, insuring a long cleft of 3 to 4 µm between contacting cells (micrograph with the green contour in Fig. 5E). Hence, despite the observation of two different junction morphologies in the static assay, the distribution of the thickness of paracellular junctions at rest was relatively homogeneous in the range of 2 to 5 µm with an average of 3.2 +/- 0.35 µm (histogram in Fig. 5E). In the pressure assay, a similar classification could be proposed in between the clusters of cells and the thin regions of the tissue. In the cell clusters, we noted that the change in cellular morphology was associated to the reorganization of nuclei, which were no longer elongated along the collagen gel scaffold (*e*.*g*., compare the panels outlined in red in Fig. 5E and 5F). The paracellular junctions, which remained aligned with the basal to apical direction (micrograph with the red contour in Fig. 5F), thickened in the range of 4 to 6 µm with an average of 5.3 µm. Conversely, in the thin regions of the tissue (micrograph with a green contour in Fig. 5F), paracellular junctions appeared to thin down with an average thickness of 1.6 µm. The resulting distribution of junction thickness in Fig. 5F was bi-modal due to these marked morphological differences. Focusing on the radius of the paracellular junctions, we did not detect the wide openings of 179 nm that were predicted by the analysis of the flux in the static *vs*. pressure assays. Our data rather hinted to the thinning of the barrier cleft as a consequence of intraluminal pressure, which we aimed to integrate in a model of paracellular transport.

### Data integration with the deformable monopore model (DMM)

We describe the DMM to recapitulate our paracellular transport data recorded in the static and pressure assay. This model assumes that (i) the pore radius is constant and equal to 24 nm in both assays (Fig. 5G-H), and (ii) the paracellular junction thickness is reduced under intraluminal pressure (Fig. 5H). The latter hypothesis implies that the diffusive permeability, which is inversely proportional to the barrier thickness according to Eq. (7), increases in the pressure assay in comparison to the static assay (denoted 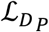 in Eq. (10)). This consequence of the deformation of the tissue changes the ratio of the diffusive to convective flux across the barrier *J_p_*/*J_D_*, as reformulated in Eq. (10). The permeation velocity to the diffusive permeability *V*_0_/*ℒ_D_* is only dependent on the paracellular pore radius (see Eq. (9)), allowing us to determine its value of 7.3×10^−3^ for an intraluminal pressure of 100 Pa. Knowing that *J_p_*/*J_D_* is 1.35 +/- 0.43, we can then fit the ratio of the diffusive permeability in the pressure to static assay 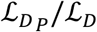 of 1.9 +/- 0.3 The DMM hence indicates that the junction thickness is divided by 1.9 upon application of intraluminal pressure. This value is in excellent agreement with electron microscopic data, which shows a reduction of the pore thickness by a factor of ∼2 from 3.2 to 1.6 µm in the thin regions of the tissue. We note that the DMM predicts that the ratio 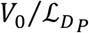 of the permeation velocity to the diffusive permeability in the pressure assay is lower than 10^−2^. Paracellular transport is thus dominated by diffusion even with intraluminal pressure, and the onset of cross-barrier flux is explained by the thinning of paracellular junctions, which is associated to an onset in diffusive permeability. Interestingly, this suggestion is borne out by the fact that the augmentation of flux across the barrier *J_p_*/*J_D_* is reduced from 1.35 to 0.67 in live *vs*. fixed samples. The fixation process indeed reduces MV deformation from 25 to 5%, and impedes the remodeling of the tissue under intraluminal pressure that is associated to the onset in diffusive permeability.

## Conclusion

We provide technologies and analytical methods to characterize molecular transport across endothelial layers in 3D microvessel structures. We prove that intraluminal pressure increases cross-barrier flux, and explain it with the DMM, which speculates that the diffusive permeability increases under pressure as a consequence of the remodeling of the endothelial tissue and the thinning of paracellular junctions. Using this model, we can specify the structural and functional properties of MV barriers (Table 1). Given the pore size *r_p_* and cleft thickness *δ* from the static assay, we deduce the pore density from Eq. (7) of ∼1 pore/μm^−2^. This estimate is in excellent agreement with the measurements in mesenteric capillaries^7^. The resulting porosity, which represents the surface fraction of pores over the MV contour, is 0.2%, showing that endothelial barriers offer little room to the passage of diffusing tracers. The permeation properties of the tissue can also be derived from Eq. (8), yielding *κ* of 10^−19^ m^2^. This value, which compares to that of impermeable shale sedimentary rocks^44^, shows that MV constitute impermeable tissues, which nevertheless enable the passage by diffusion of molecules of ∼10 nm, a size range that matches that of most plasma proteins^45^. Finally, we suggest that paracellular transport across endothelial tissues can be controlled by their deformation without any change in adherens junction protein expression. It is thus tentative to speculate that the stiffening of the supporting matrix, which is an age-related phenotype^46^, can restrict the deformation of the barrier, reduce exchange through the endothelium, and contribute to unbalanced homeostasis in aged tissues.

**Table 1:** Overview on MV properties obtained from the DMM for an intraluminal pressure of 100 Pa.

| Pore size<br>$r_p$ (nm) | Paracellular<br>barrier thickness<br>$\delta$ ($\mu\text{m}$) | Pore density<br>$n$ ($\mu\text{m}^{-2}$) | Porosity<br>$\varphi$ (%) | Permeability<br>$\kappa$ ($10^{-19} \text{ m}^2$) |
| --- | --- | --- | --- | --- |
| 24 +/- 3 | 3.2 +/- 0.4 | 0.85 +/- 0.31 | 0.16 +/- 0.05 | 1.2 +/- 0.6 |

## Materials and Methods

### Cell culture and reagents

All chemicals were purchased from Sigma Aldrich, unless mentioned. Primary human umbilical vein endothelial cells (HUVEC; Lonza, Basel, Switzerland) were cultured in Endothelial Cell Growth Medium-2 BulletKit (EGM-2; Lonza) and used between passages 4 to 7.

### MV fabrication

Microvessel (MV) were fabricated in polydimethylsiloxane (PDMS)-based chips (25 mm × 25 mm × 5 mm: width × length × height), as previously described^24^. The protocol includes an additional PDMS-collagen cross-linking step to avoid leaks at the PDMS/collagen interface during diffusion and pressure assays. The protocol started by O2 plasma treatment of PDMS chips and acupuncture needles of 200 μm (No. 08, J type; Seirin, Shizuoka, Japan) for one minute (basic plasma cleaner; Harrick Plasma, Ithaca, NY, USA). The PDMS chips and needles were then placed together in a vacuum reactor with 100 μL of aminopropyl-triethoxysilane, and left at 0.1 mbar and room temperature for 30 minutes. Needles were then soaked in 1% (w/v) bovine serum albumin, dried, and sterilized by UV-light exposure. The chips were treated with 50 μL of 2.5% glutaraldehyde (GA) for 1 minute, then thoroughly rinsed with water, and dried. The collagen solution was subsequently prepared on ice by mixing Cellmatrix® Type I-A collagen solution (Nitta Gelatin, Japan), 10× Hanks’ buffer, and 10× collagen buffer (volume ratio 8:1:1) following manufacturer’s protocol (final collagen concentration: 2.4 mg/mL). We poured 30 μL of this ice-cold collagen solution into the chip, and inserted the coated needle. The devices were incubated at 37°C for 40 min to induce collagen reticulation, and the needles were withdrawn to form a hollow channel. The chips were left in PBS at least overnight before cell seeding, and the holes for needle incorporation were sealed with unreticulated PDMS to prevent leaks during the diffusion and pressure assays.

Just prior to loading in the chips, HUVEC cells were harvested and resuspended in the medium supplemented with 3% (m/v) dextran (500 kDa) at a density of 10^7^ cells/mL. 50,000 cells were loaded at each opening of the channel, and let to attach to the collagen scaffold at 37°C for 10 minutes. This operation was repeated one time to insure a high degree of coverage (over 80%) inside the chip. Warm medium was finally added, and MV were cultured at 37°C until use two days after fabrication.

### Fixation, immunostaining and electron microscopy

For structural analysis, MV were fixed with 4% paraformaldehyde (PFA) or 2.5% GA at 37°C for 60 min, and then thoroughly rinsed with PBS. PFA fixed samples were analyzed by the diffusion/pressure assays. They were also used for immunostaining starting with permeabilization with 0.5% Triton X-100 for 10 min. Blocking with 1% BSA was performed overnight at 4°C. Cells were incubated overnight at 4°C with the primary antibody against VE-Cad (rabbit mAb, D87F2, Cell Signaling Technology, 1:200) diluted in blocking solution. After washing, cells were incubated for 2 h with the secondary antibody (1:400) and Alexa Fluor-488 Phalloidin (1:400). Labeled samples were washed and stored at 4°C until imaging. Confocal images were captured with the LSM 700 confocal microscope (Carl Zeiss) equipped with a 40× water immersion objective (numerical aperture (NA) of 1.2). We used a pinhole of 1 Airy unit for the three lasers of 488 and 555 nm, and set the increment between confocal stacks to 1.0 μm.

For electron microscopy samples were fixed with 2.5% glutaraldehyde-0.1 M phosphate buffer (pH 7.4). They were postfixed with Osmium Tetroxide, dehydrated in a series of graded ethanol, embedded in epoxy resin Epon 812, then cut into ultrathin sections. The sections were stained with uranyl acetate and lead citrate, and examined using transmission electronic microscope (TEM; JEM-1011, JEOL, Japan).

### Setting up the diffusion and pressure assays

MV were placed on an aluminum support of 30 × 60 mm^2^ with a set of tapped holes to tightly hold 3D printed reservoirs fabricated by stereolithography (Expert Material Series, NSS, Japan). Fluorescence redistribution experiments were conducted by confocal microscopy setting the optical section to 12 μm (1 Airy unit of a 10× air objective (NA=0.4), i.e., a size smaller than MV diameter. The inter-frame time interval was set to 7.7 s with an image size of 512×512 pixels^2^, equivalently 1.28×1.28 mm^2^. We used the 4 kDa dextran coupled to FITC and the 70 kDa dextran coupled to rhodamine-B. They were loaded inside the lumen at a concentration of 1.2 mg/mL in culture medium. Note that the macromolecules used in the static assay were rinsed with culture medium and the samples were placed in the incubator for ∼30 minutes before running the pressure assay. The diffusion coefficient *D*_0_ of these macromolecules was 244 +/- 25 and 62 +/- 6 µm^2^/s in collagen gels, respectively (Supplementary Fig. S2). The ratio between these measurements was expectedly consistent with the ratio of their hydraulic radii *r_dye_* of 1.5, and 6.2 nm^47,48^.

Image analysis was performed with ImageJ. The radius of the tube *r*_0_ was obtained by applying a Canny edge detection filter (FeatureJ Edges), and by fitting the pixel intensity profile perpendicularly to the tissue with an error function (Supplementary Fig. S1). Final data were analyzed and fitted with Igor Pro. Average data were reported with standard errors.

### COMSOL simulations

Simulations were run with COMSOL Multiphysics 6.0. We used the Transport of Diluted Species in Porous Media module based on the tortuosity model with a tortuosity equal to unity, and the Darcy law module to compute the flow velocity field. We set the tube radius *r*_0_ to 100 μm and placed it 280 μm above the glass coverslip. The barrier thickness *δ* was set to 3 μm. We integrated the simulation data over a field of 1.2 mm centered around the tube. Three distinct domains were defined: (i) the collagen matrix with a porosity of 0.95^49^, (ii) the central lumen free of collagen, and (iii) the cell barrier as characterized by a tunable diffusive permeability and permeation velocity. The permeability of collagen gels was set to 5×10^−14^ m^2 50,51^. Boundary conditions consisted of no flux at the edges of the gel expect for an open boundary at the top surface and tube interface. The initial dye concentration was set to 0.025 mol/m^3^ in the lumen and to null in the collagen matrix. The pressure was set to 100 Pa in the lumen and to null at the top interface of the gel.

## Acknowledgements

JC acknowledges the JSPS for postdoctoral fellowship. DA acknowledges the WINGS-QSTEP program for doctoral fellowship. This research was partly supported by the Grant-in-Aid for JSPS Fellows (20F20806), AMED P-CREATE (JP18cm0106239h0001), JSPS Core-to-Core Program (JPJSCCA20190006), AMED-CREST (22gm1510009). The authors thank the LIMMS (CNRS-Institute of Industrial Science, University of Tokyo) for financial support, and Eri Otsuka, Tadaaki Nakajima, and Takanori Sano for technical assistance and critical reading of the manuscript. The authors also thank Sachie Matsubara (Laboratory for Electron Microscopy, Kyorin University School of Medicine) for technical assistance with electron macroscopy.

## SUPPLEMENTARY MATERIAL

**Supplementary Fig. S1:**
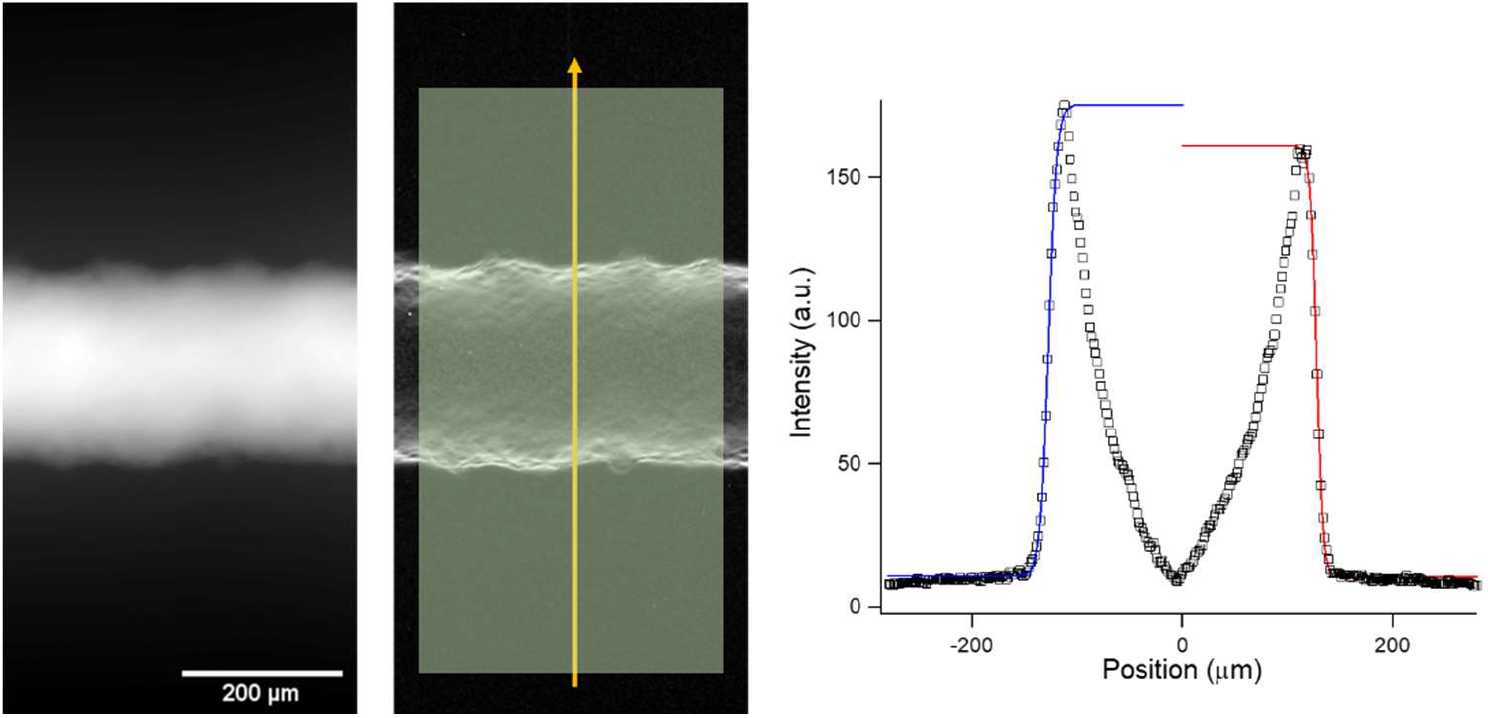
Method to determine the MV radius *r*_0_. Starting from the fluorescence intensity signal averaged over the entire stack (left panel), we used a Cany edge detection filter in Imagej (middle panel), extracted the intensity plot across the microvessel (yellow arrow in the middle panel), and fitted the position of vessel edges with error functions that are indicated with blue and red curves on the graph in the right.

**Supplementary Fig. S2:**
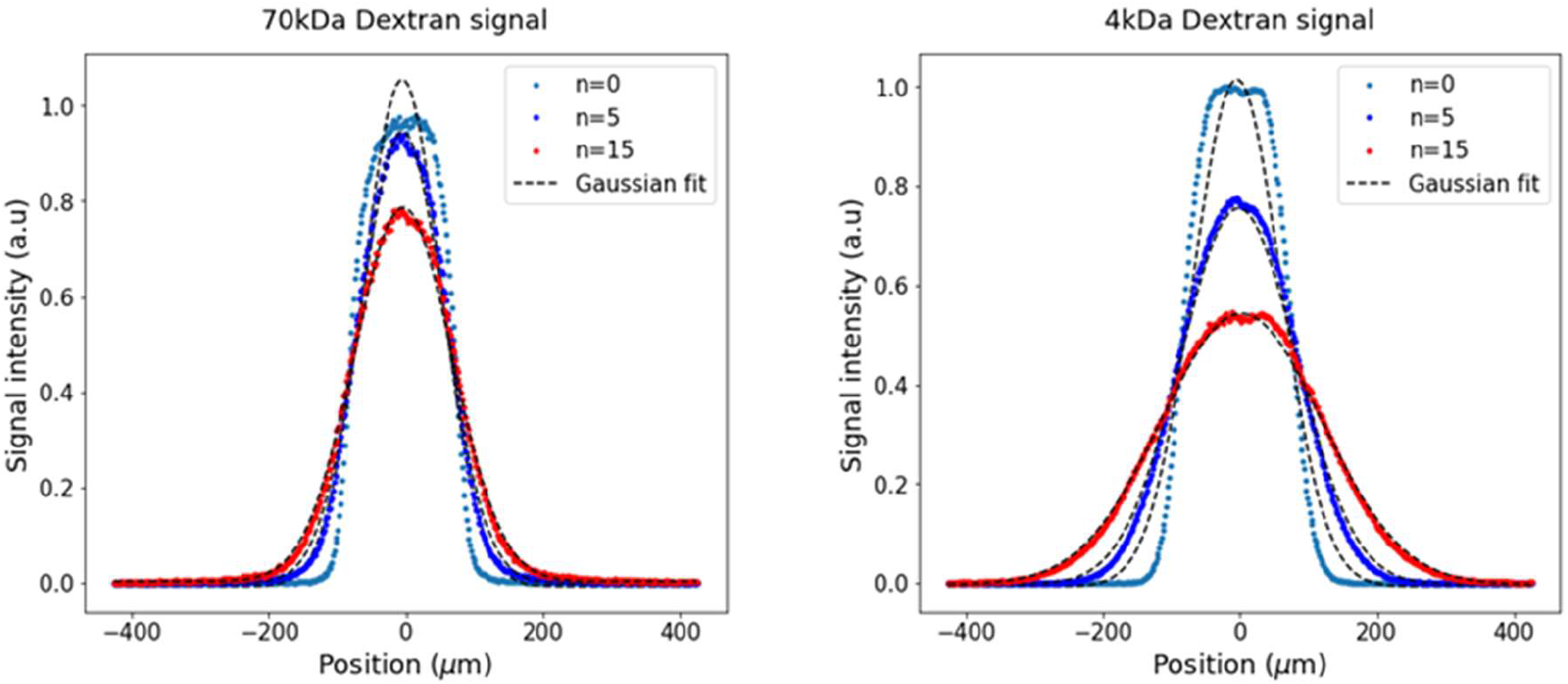
Representative fluorescence intensity distribution over time for FITC- and rhodamine-dextran diffusing in collagen gel. Using the static assay on a collagen lumen with no cells, we co-inject the 70 kDa and 4 kDa dextran, and monitor fluorescence redistribution over time with an inter-frame interval of 1.9 s and image sizes of 512*100 pixels^2^. Note that n correspond to the image number in the time series. Data are fitted with Gaussian functions (dashed lines) to extract the full width at half maximum over time. The width increases linearly with time and proportionally with the diffusion coefficient. This method has been calibrated and validated with COMSOL simulations.

**Supplementary Fig. S3:**
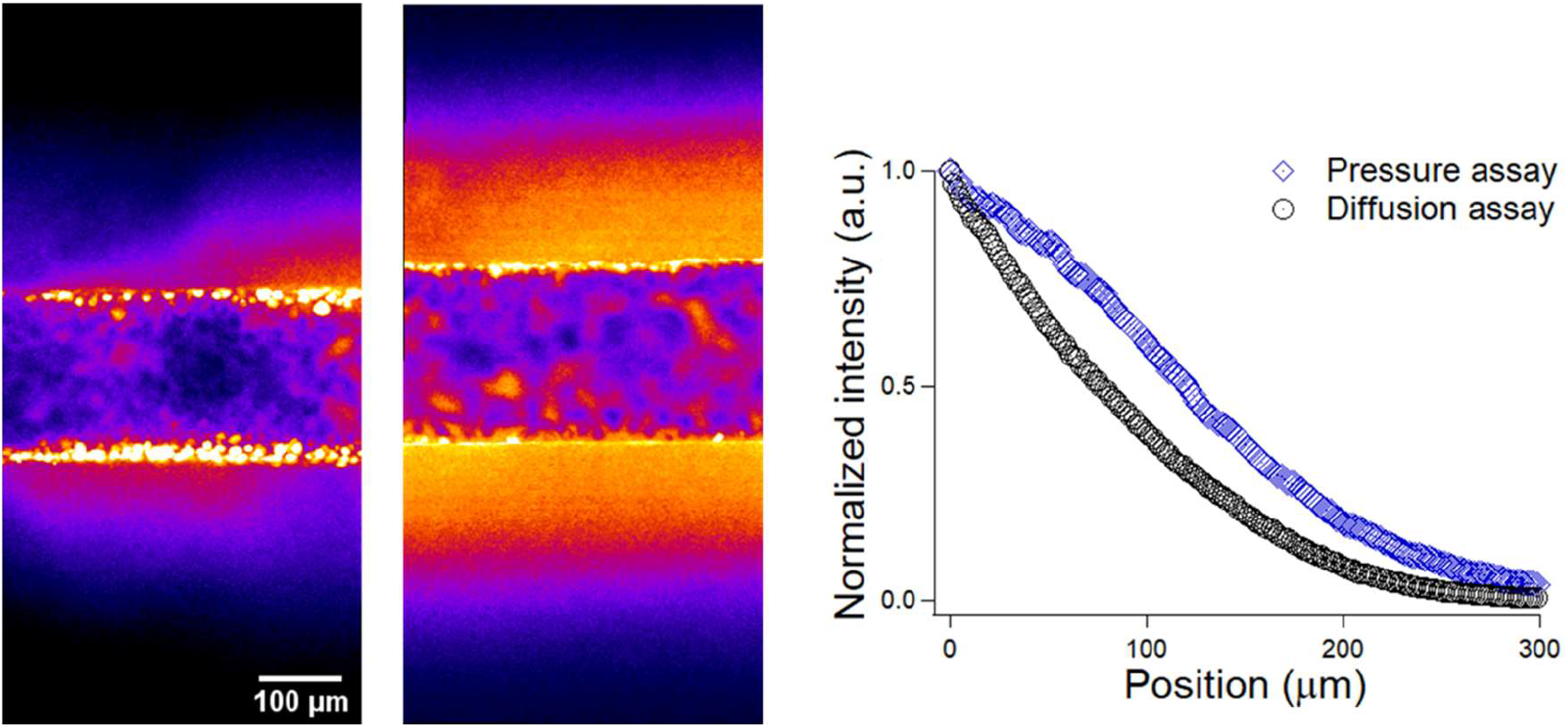
Concentration profiles after 30 s in leaky barriers. Fluorescence micrographs are obtained with the static and pressure assays (left and middle panels, respectively). The graph in the right shows the respective normalized intensity profiles in the collagen gel, as indicated in the legend.

## References

1 J. Mullin, N. Agostino, E. Rendonhuerta and J. Thornton, Drug Discov. Today, 2005, 10, 395–408.

2 G. Bazzoni and E. Dejana, Physiol. Rev., 2004, 84, 869–901.

3 M. Z. Gładysz, M. Stevanoska, M. K. Włodarczyk-Biegun and A. Nagelkerke, Adv. Drug Deliv. Rev., 2022, 184, 114183.

4 Y. Komarova and A. B. Malik, Annu. Rev. Physiol., 2010, 72, 463–493.

5 R. D. Minshall, C. Tiruppathi, S. M. Vogel and A. B. Malik, Histochem. Cell Biol., 2002, 117, 105–112.

6 K. Y. Y. Fung, G. D. Fairn and W. L. Lee, Traffic, 2018, 19, 5–18.

7 B. Rippe and B. Haraldsson, Physiol. Rev., 1994, 74, 163–219.

8 Y. Wallez and P. Huber, Biochim. Biophys. Acta BBA - Biomembr., 2008, 1778, 794–809.

9 L. Claesson-Welsh, E. Dejana and D. M. McDonald, Trends Mol. Med., 2021, 27, 314–331.

10 S. Sindhwani, A. M. Syed, J. Ngai, B. R. Kingston, L. Maiorino, J. Rothschild, P. MacMillan, Y. Zhang, N. U. Rajesh, T. Hoang, J. L. Y. Wu, S. Wilhelm, A. Zilman, S. Gadde, A. Sulaiman, B. Ouyang, Z. Lin, L. Wang, M. Egeblad and W. C. W. Chan, Nat. Mater., 2020, 19, 566–575.

11 B. R. Kingston, Z. P. Lin, B. Ouyang, P. MacMillan, J. Ngai, A. M. Syed, S. Sindhwani and W. C. W. Chan, ACS Nano, 2021, 15, 14080–14094.

12 D. Knowland, A. Arac, K. J. Sekiguchi, M. Hsu, S. E. Lutz, J. Perrino, G. K. Steinberg, B. A. Barres, A. Nimmerjahn and D. Agalliu, Neuron, 2014, 82, 603–617.

13 U. Kniesel and H. Wolburg, Cell. Mol. Neurobiol., 2000, 20, 57–76.

14 P. Artursson, J. Pharm. Sci., 1990, 79, 476–482.

15 C. Gupta, A. Chauhan and S. P. Srinivas, Pharm. Res., 2012, 29, 3325–3334.

16 C. C. Michel and F. E. Curry, Physiol. Rev., 1999, 79, 703–761.

17 N. Ferrell, R. R. Desai, A. J. Fleischman, S. Roy, H. D. Humes and W. H. Fissell, Biotechnol. Bioeng., 2010, 107, 707–716.

18 A. Martinez-Palomo, I. Meza, G. Beaty and M. Cereijido, J. Cell Biol., 1980, 87, 736–745.

19 M. Cereijido, E. Robbins, W. Dolan, C. Rotunno and D. Sabatini, J. Cell Biol., 1978, 77, 853–880.

20 E. Biganzoli, L. A. Cavenaghi, R. Rossi, M. C. Brunati and M. L. Nolli, Il Farm., 1999, 54, 594–599.

21 21 E. W. K. Young, M. W. L. Watson, S. Srigunapalan, A. R. Wheeler and C. A. Simmons, Anal. Chem., 2010, 82, 808–816.

22 S. M. Albelda, P. M. Sampson, F. R. Haselton, J. M. McNiff, S. N. Mueller, S. K. Williams, A. P. Fishman and E. M. Levine, J. Appl. Physiol., 1988, 64, 308–322.

23 A. Thomas, S. Wang, S. Sohrabi, C. Orr, R. He, W. Shi and Y. Liu, Biomicrofluidics, 2017, 11, 024102.

24 E. Lee, H. Takahashi, J. Pauty, M. Kobayashi, K. Kato, M. Kabara, J. Kawabe and Y. T. Matsunaga, J. Mater. Chem. B, 2018, 6, 1085–1094.

25 J. Pauty, R. Usuba, I. G. Cheng, L. Hespel, H. Takahashi, K. Kato, M. Kobayashi, H. Nakajima, E. Lee and F. Yger, EBioMedicine, 2018, 27, 225–236.

26 R. Bednarek, Methods Protoc., 2022, 5, 17.

27 C. A. Dessalles, C. Ramón-Lozano, A. Babataheri and A. I. Barakat, Biofabrication,, DOI:10.1088/1758-5090/ac2baa.

28 T. J. Lohela, T. O. Lilius and M. Nedergaard, Nat. Rev. Drug Discov., 2022, 21, 763–779.

29 T. Nakajima, K. Sasaki, A. Yamamori, K. Sakurai, K. Miyata, T. Watanabe and Y. T. Matsunaga, Biomater. Sci., 2020, 8, 5615–5627.

30 J. Pauty, R. Usuba, H. Takahashi, J. Suehiro, K. Fujisawa, K. Yano, T. Nishizawa and Y. T. Matsunaga, Nanotheranostics, 2017, 1, 103.

31 K. Mari Hämäläinen, K. Kontturi, S. Auriola, Lasse Murtomäki and A. Urtti, J. Controlled Release, 1997, 49, 97–104.

32 P. Dechadilok and W. M. Deen, Ind. Eng. Chem. Res., 2006, 45, 6953–6959.

33 J. L. Snyder, A. Clark, D. Z. Fang, T. R. Gaborski, C. C. Striemer, P. M. Fauchet and J. L. McGrath, J. Membr. Sci., 2011, 369, 119–129.

34 J. Bear, Dynamics of fluids in porous media, Dover, New York, 1988.

35 N. Nishiyama and T. Yokoyama, J. Geophys. Res. Solid Earth, 2017, 122, 6955–6971.

36 G. S. Offeddu, K. Haase, M. R. Gillrie, R. Li, O. Morozova, D. Hickman, C. G. Knutson and R. D. Kamm, Biomaterials, 2019, 212, 115–125.

37 E. M. Renkin, in Endothelial Cell Biology in Health and Disease, eds. N. Simionescu and M. Simionescu, Springer US, Boston, MA, 1988, pp. 51–68.

38 M. Hahn, T. Heubach, A. Steins and M. Jünger, J. Invest. Dermatol., 1998, 110, 982–985.

39 Y. Tian, G. Gawlak, J. J. O’Donnell, A. A. Birukova and K. G. Birukov, J. Biol. Chem., 2016, 291, 10032–10045.

40 A. A. Birukova, N. Moldobaeva, J. Xing and K. G. Birukov, Am. J. Physiol.-Lung Cell. Mol. Physiol., 2008, 295, L612–L623.

41 G. Punshon, D. Vara, K. Sales, A. Kidane, H. Salacinski and A. Seifalian, Biomaterials, 2005, 26, 6271–6279.

42 P. M. Davidson and B. Cadot, Trends Cell Biol., 2021, 31, 211–223.

43 K. Burridge and E. S. Wittchen, J. Cell Biol., 2013, 200, 9–19.

44 R. A. Freeze and J. A. Cherry, Groundwater, Prentice-Hall, Englewood Cliffs, N.J, 1979.

45 H. P. Erickson, Biol. Proced. Online, 2009, 11, 32–51.

46 M. J. Sherratt, Adv. Wound Care, 2013, 2, 11–17.

47 M. B. Mustafa, D. L. Tipton, M. D. Barkley, P. S. Russo and F. D. Blum, Macromolecules, 1993, 26, 370–378.

48 A. Wolde-Kidan, A. Herrmann, A. Prause, M. Gradzielski, R. Haag, S. Block and R. R. Netz, Biophys. J., 2021, 120, 463–475.

49 T. Fischer, A. Hayn and C. T. Mierke, Sci. Rep., 2019, 9, 8352.

50 E. Gentleman, E. A. Nauman, K. C. Dee and G. A. Livesay, Tissue Eng., 2004, 10, 421–427.

51 V. Serpooshan, M. Julien, O. Nguyen, H. Wang, A. Li, N. Muja, J. E. Henderson and S. N. Nazhat, Acta Biomater., 2010, 6, 3978–3987.

